# Engineering a bifunctional alfa and beta hydrolase from a GH1 beta-glycosidase

**DOI:** 10.64898/2026.03.19.712844

**Authors:** Felipe A.M. Otsuka

## Abstract

Glycoside hydrolases (GHs) play central roles in carbohydrate metabolism and are widely exploited for industrial and biomedical applications. However, they are often not optimal for applications due to their constrained function and strict stereochemical specificity, necessitating the discovery and optimization of distinct enzymes for each glycosidic configuration. Members of glycoside hydrolase family 1 (GH1) are archetypal retaining β-glycosidases, while α-specific activity is rare within this family. Here, I demonstrate that a retaining GH1 enzyme can be engineered to hydrolyze both β- and α-configured substrates without altering its canonical catalytic residues. Using a well-characterized β-glycosidase and computational protein design strategies targeting second-shell residues surrounding the active site, a bifunctional β-/α-glycosidase containing 45 mutations was generated. The engineered variant acquired the ability to hydrolyze the α-configured substrate 4-nitrophenyl-α-D-glucopyranoside while retaining activity toward the originals β-substrates, with reduced catalytic efficiency and thermostability. Structural modeling and docking analyses reveal that the engineered enzyme preserves the original fold and accommodates substrates within the catalytic pocket in a similar manner to the wild type. These findings provide direct evidence that stereochemical constraint in retaining GH is more flexible than previously appreciated and can be modulated through targeted engineering.

## Introduction

Glycoside hydrolases (GH) are ubiquitous enzymes that catalyze the hydrolysis of glycosidic bonds in oligo- and polysaccharides, glycoconjugates, and glycolipids, thereby shaping carbon flux, host–microbe interactions and building cellular architecture across all domains of life^1^. The biological relevance of glycans together with GHs justify their use in industrial and biomedical applications, such as lignocellulosic biomass saccharification for bioethanol production, and the remodeling of cell-surface glycans for therapeutic purposes^2–4^.

Glycosidases are classified in the Carbohydrate-Active enZymes (CAZy) database^5^ to facilitate annotation of GHs according to sequence, structure and function. Members of glycoside hydrolase family 1 (GH1) are well-characterized enzymes belonging to clan GH-A and adopt the canonical (β/α)_8_-barrel fold. GH1 enzymes are retaining β-glycosidases (**Figure 1C**) that operate via the Koshland double-displacement mechanism, in which a catalytic nucleophile and a general acid/base residue (typically glutamate) mediate formation and breakdown of a covalent glycosyl– enzyme intermediate, resulting in the retention of configuration at the anomeric center^6^. A notable feature of the GH1 family is its pronounced stereochemical bias toward β-configured substrates (*e*.*g*., β-glucosidase and β-galactosidase), whereas activities toward α-linked substrates are rare (**Figure 1A-B**), such as α-L-arabinopyranosidase and exo-α-neuraminidase (sialidase). This distribution indicates that the GH1 active-site architecture is strongly optimized for β-configured substrates.

**Figure 1.**
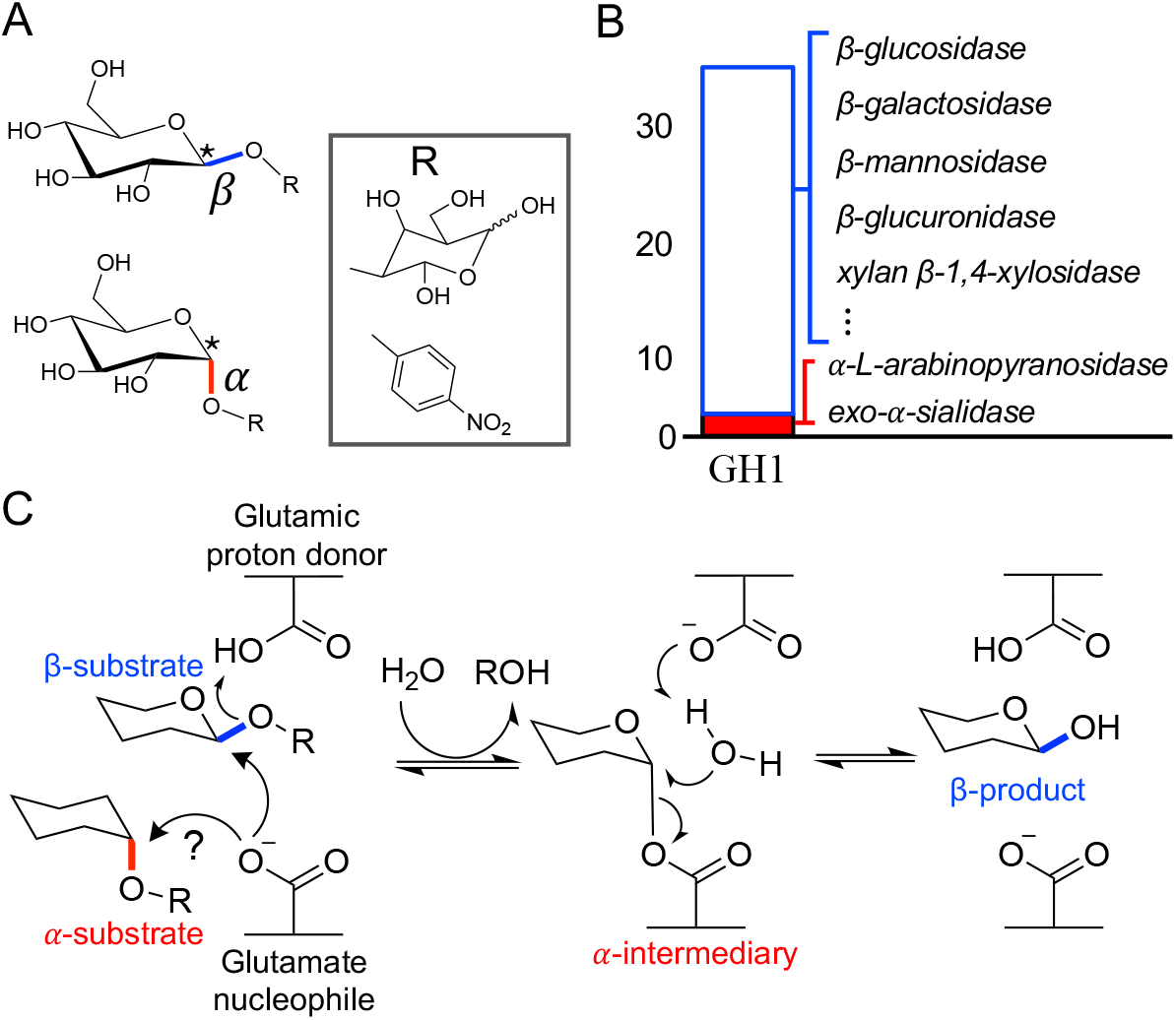
Stereospecificity bias in glycoside hydrolase family 1 (GH1). (**A**) Schematic representation of β- and α-anomeric configurations at the anomeric carbon of a pyranose sugar. The “R” group can be another sugar or a 4-nitrophenol group (Pnp) (**B**) Counts of reported GH1 activities toward β-(empty blue bar) and α-(red fill bar) substrates, based on curated CAZyme annotations. A total of 32 distinct β-specific GH1 activities have been reported, compared with only 2 α-specific activities. Examples of GH1 enzymes with β-substrate specificity, including β-glucosidase (EC 3.2.1.21), β-galactosidase (EC 3.2.1.23), β-mannosidase (EC 3.2.1.25), and many others. (**C**) Retaining hydrolysis mechanism of GH1 enzymes. A β-substrate (blue bond) is processed to yield a β-product via a glycosyl α-intermediate. In contrast, α-substrates processing in GH1 is uncertain due to the optimal positioning of the catalytic residues in the active site for β-substrates.

Reports of GHs capable of catalyzing both β- and α-substrates for the same glycosidic linkage are scarce, and to date no targeted engineering strategy has been developed to address this matter. Importantly, the retaining double-displacement mechanism itself does not inherently preclude specialization toward either β- or α-configuration, as evidenced by retaining glycosidases that are naturally specific for α-configured substrates^7,8^. This observation raises a fundamental mechanistic question: can a single retaining active site accommodate and catalyze hydrolysis of opposing anomeric configurations for the same substrate (**Figure 1C**)? Moreover, the immense structural and stereochemical diversity of glycans hinders the readily utilization of GHs, largely because their limited enzymatic flexibility toward substrates^9^. Demonstrating a glycosidase activity toward the opposite anomeric configuration would constitute a proof of concept for expanding GH1 functionality.

Enzyme engineering offers a powerful approach to explore such non-native functions while providing insight into fundamental aspects of catalytic mechanism^10^. In this work, I address these challenges by engineering a native β-specific GH to hydrolyze its cognate α-configured substrate. I used a well-characterized GH1 β-glycosidase from *Spodoptera frugiperda* (Sfβgly)^11^, an enzyme with broad activity toward β-substrates and notable robustness to mutation while retaining thermostability and catalytic efficiency^12,13^. Using computational protein design to explore the functional sequence space of second-shell residues surrounding the active site, I generated a glycosidase variant capable of hydrolyzing both β-configured substrates and the α-configured analog 4-nitrophenyl-α-D-glucopyranoside. Structural modeling and docking analysis indicate that the engineered variant preserves the original fold while accommodating substrates within the catalytic pocket in a similar manner to the WT Sfβgly. Together, these results establish that a retaining GH1 enzyme can be engineered to process opposing anomeric configurations of the same substrate without altering the canonical catalytic residues. Moreover, it reveals an unexpected flexibility in the stereochemical constraints governing glycosidic bond hydrolysis.

## Results

### Engineering a bifunctional α- and β-glycosidase

To explore the possibility of introducing new substrate into a native β-GH, I used Rosetta^14^ to minimize the X-ray structure and PROSS^15^, a designing algorithm that suggest mutations to improve protein solubility and stability in *E. coli*. Amino acid positions directly contacting the substrate in the catalytic site (first-shell residues) and at the dimerization interface were restricted from mutation. The resulting *in-silico* variant, designated PROSS_design3, contained 41 mutations relative to the WT Sfβgly (Fig. S1). This PROSS-optimized sequence was then used as input for FuncLib^16^, a design tool that diversifies selected residues surrounding the active site. Mutations in the vicinity of the catalytic site were avoided because they are often detrimental activity; however, second-shell positions can influence the coordination of the catalytic residues^17^ and may enable accommodation of alternative substrates. Among the several designs generated, one cluster reached a tolerable sequence space. The five top-scoring models were visually inspected, and one sequence was selected for experimental validation (**Figure 2A-B**, Fig. S1-S3). The final variant contained, in addition to mutations by PROSS, four more mutations relative to the WT: Y38W, W143F, L372M and S445C.

**Figure 2.**
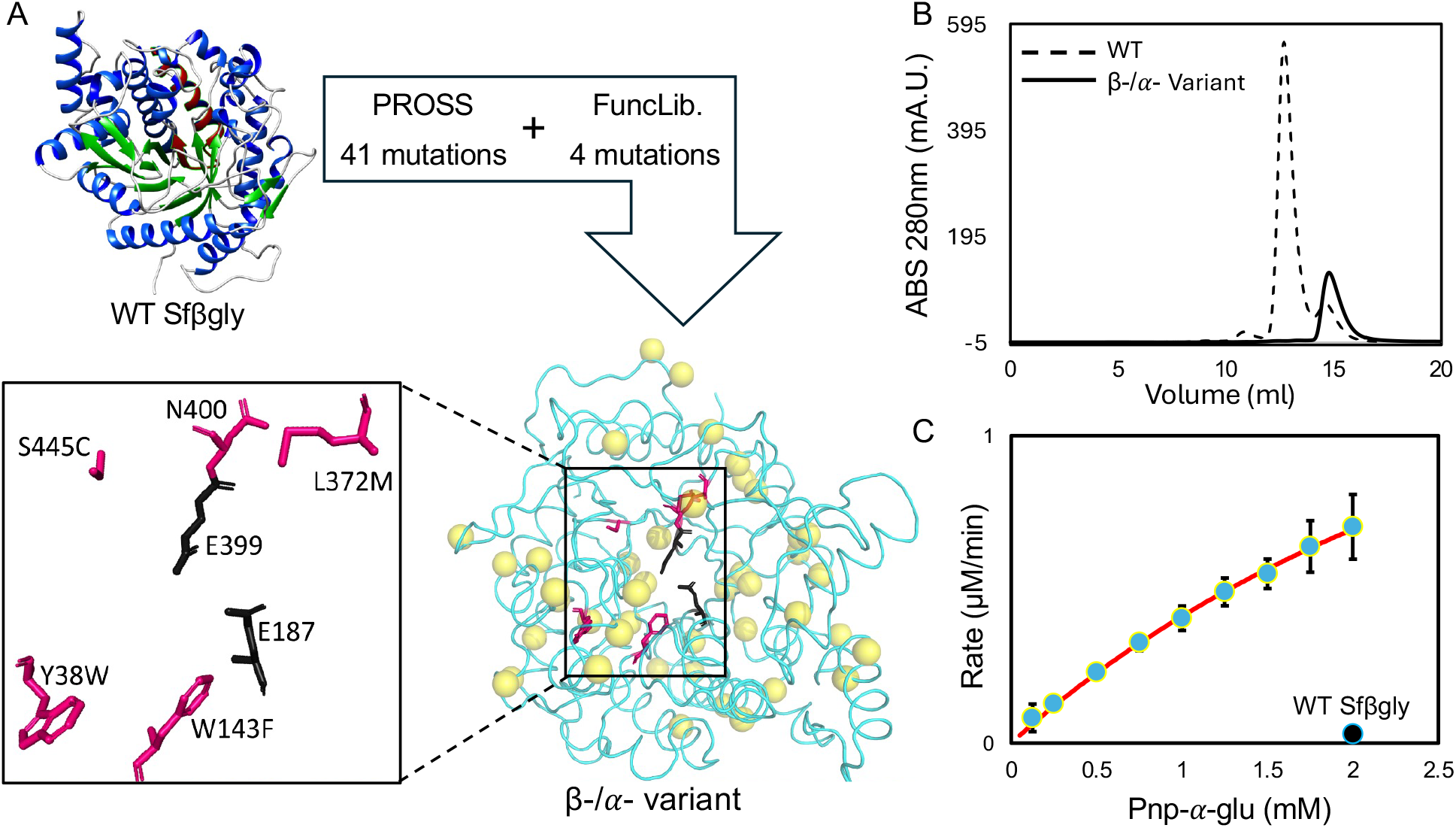
Engineering of a β-/α-glycosidase using combined protein design pipeline. (**A**) The workflow started with the WT Sfβgly structure and submitted to PROSS for generating a more soluble and stable mutant, the generated sequence was directly used in FuncLib focusing on five residues in the second shell of the catalytic pocket. The predicted structure of the variant shows mutations made by PROSS in yellow spheres, and the positions selected to FuncLib in purple. Catalitic residues E187 and E399 are in black. (**B**) Size-exclusion chromatography comparison of the WT Sfβgly (dashed line) and the β-/α-variant (solid line). (**C**) Enzymatic assay of the β-/α-variant (blue dots) fitted with a Michaelis-Menten equation for 4h using Pnp-α-glu as substrates, a black dot indicates the measurement of the WT Sfβgly under the same assay for the highest substrate concentration. Data from two independent replicates. Sum of squared residuals from the fit is 0.0015.

The engineered variant was cloned into pET-28a(+) and expressed in *E. coli* BL21 (DE3). After purification (Fig. S2-S3), size-exclusion chromatography revealed that the variant is a monomer, whereas the WT Sfβgly display a major dimeric fraction^18^ (**Figure 2B**). This shift suggests that the mutations introduced by PROSS increased monomer solubility, even though dimer-interface residues were preserved during design; thus, the enhanced monomeric state is not attributable to disruption of the native interface. Functionally, the one-shot variant displayed clear activity toward the α-substrate Pnp-α-glu (**Figure 2C**), while the WT Sfβgly showed no detectable catalysis under parallel conditions. These findings demonstrate that a protein design strategy targeting second-shell residues can successfully convert a native β-glycosidase to catalyze an α-configured substrate.

### Kinetic and thermostability profile of β-/α-variant versus WT Sfβgly

To assess how much of the original enzymatic function was retained in the β-/α-variant, the kinetic parameters of four substrates were measured and compared with those of the WT Sfβgly (Fig. S4). As shown in **Table 1** and **Figure 3A** acquisition of α-hydrolytic activity came at a substantial cost to the hydrolysis of the native β-substrates. In particular, the catalytic efficiency toward Pnp-β-glu decreased by approximately 2.6 orders of magnitude in the variant, with both *K*_M_ and *k*_cat_ impaired. The variant remained capable of hydrolyzing the other two β-glycosidase substrates, Pnp-β-fuc with roughly one-order less efficient due to a decrease in *k*_cat_, and Pnp-β-gal, with a sixfold reduction in efficiency due to changes in both kinetic parameters. Catalytic values measured for the WT Sfβgly were consistent with previously reported data^11,19^. Collectively, these results indicate that the engineered enzyme retains its β-hydrolase activity while gaining the ability to process an α-configured substrate, demonstrating true dual specificity.

**Table 1.**
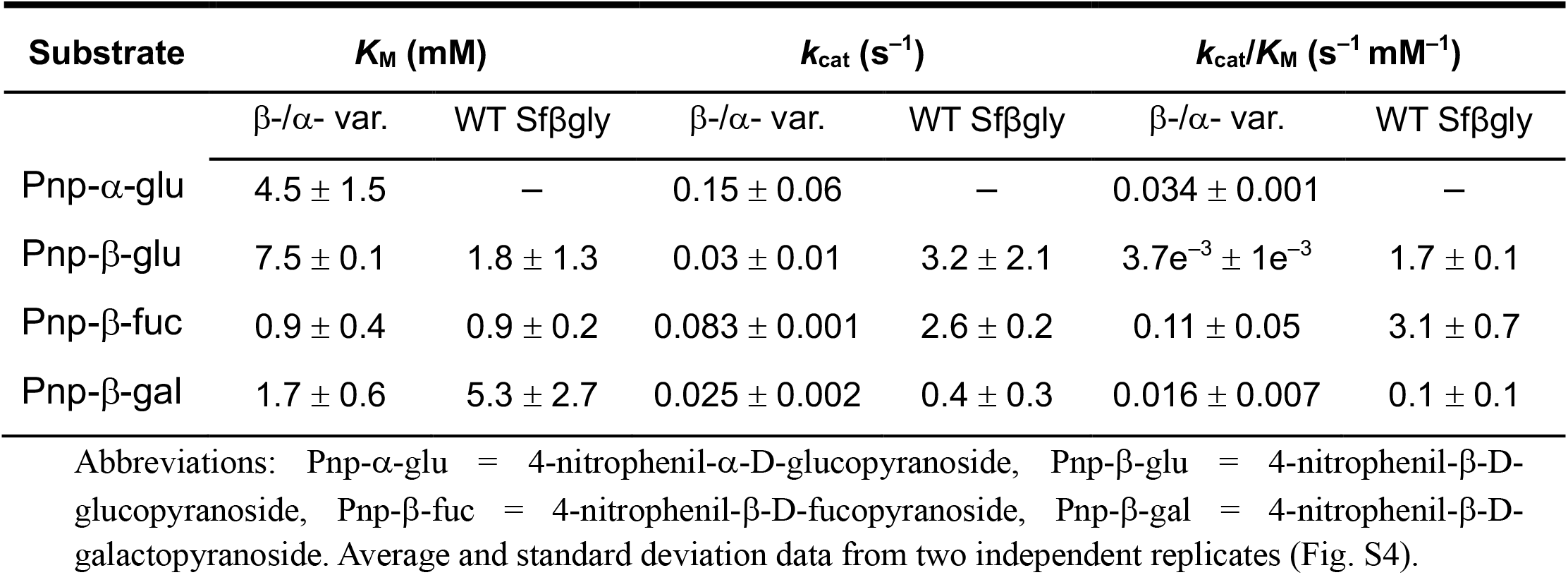
Comparison of Michaelis-Menten kinetic parameters of four different Pnp substrates.

| Substrate | $K_M$ (mM) | | $k_{cat}$ ($s^{-1}$) | | $k_{cat}/K_M$ ($s^{-1} mM^{-1}$) | |
| --- | --- | --- | --- | --- | --- | --- |
| | β-/α- var. | WT Sfβgly | β-/α- var. | WT Sfβgly | β-/α- var. | WT Sfβgly |
| Pnp-α-glu | $4.5 \pm 1.5$ | — | $0.15 \pm 0.06$ | — | $0.034 \pm 0.001$ | — |
| Pnp-β-glu | $7.5 \pm 0.1$ | $1.8 \pm 1.3$ | $0.03 \pm 0.01$ | $3.2 \pm 2.1$ | $3.7e^{-3} \pm 1e^{-3}$ | $1.7 \pm 0.1$ |
| Pnp-β-fuc | $0.9 \pm 0.4$ | $0.9 \pm 0.2$ | $0.083 \pm 0.001$ | $2.6 \pm 0.2$ | $0.11 \pm 0.05$ | $3.1 \pm 0.7$ |
| Pnp-β-gal | $1.7 \pm 0.6$ | $5.3 \pm 2.7$ | $0.025 \pm 0.002$ | $0.4 \pm 0.3$ | $0.016 \pm 0.007$ | $0.1 \pm 0.1$ |

Abbreviations: Pnp-α-glu = 4-nitrophenil- α-D-glucopyranoside, Pnp-β-glu = 4-nitrophenil- β-D-glucopyranoside, Pnp-β-fuc = 4-nitrophenil- β-D-fucopyranoside, Pnp-β-gal = 4-nitrophenil- β-D-galactopyranoside. Average and standard deviation data from two independent replicates (Fig. S4).

**Figure 3.**
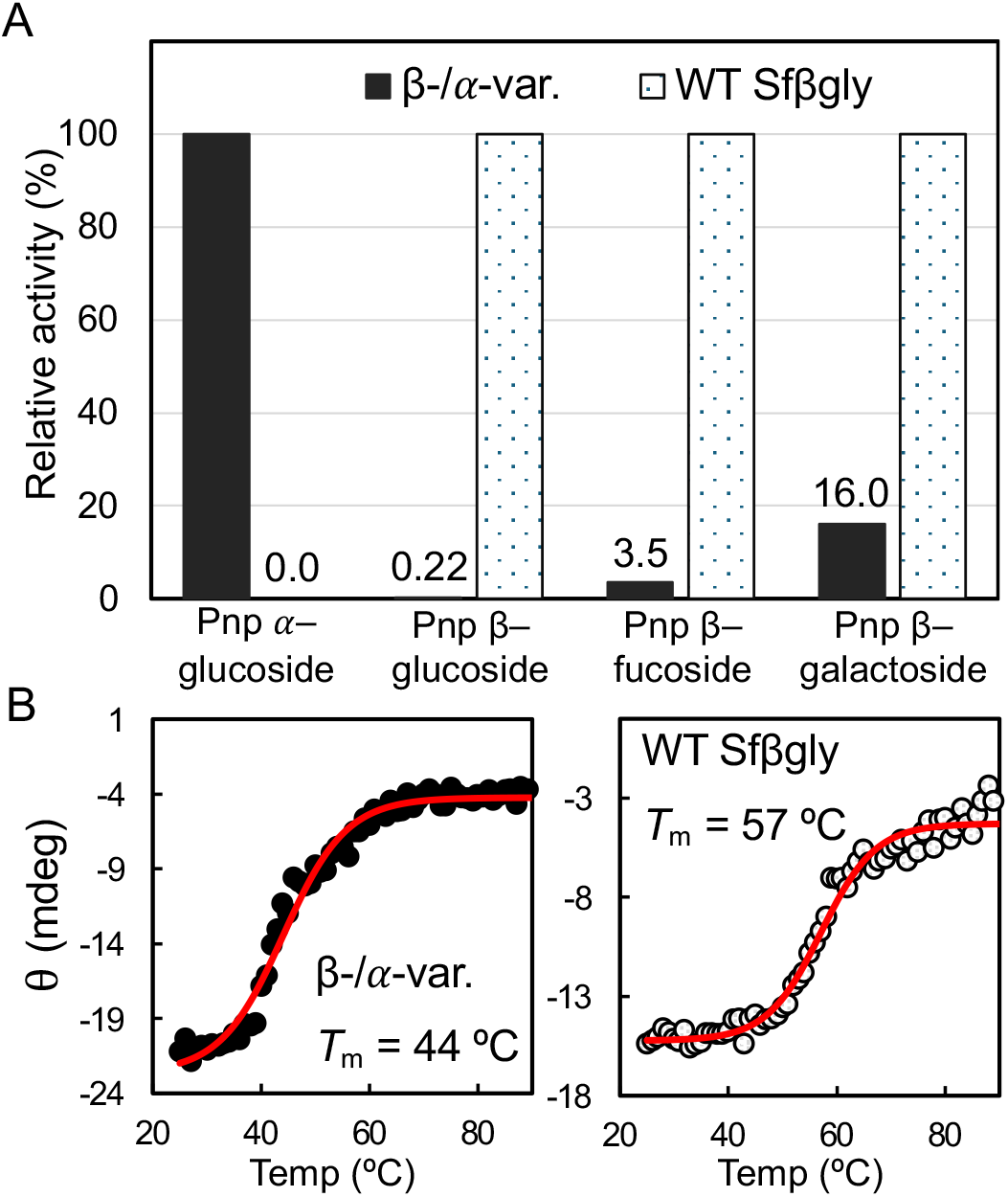
Comparison of catalytic efficiency and thermal stability between β-/α-variant and WT Sfβgly. (**A**) Relative catalytic efficiency of both enzymes toward the four tested substrates. (**B**) Thermal unfolding profiles measured by circular dichroism at 220 nm. Full CD spectrum profiles at 20 °C and 90 °C are shown in Fig. S5.

To address possible alterations in the fold from the accumulated mutations in the variant, I performed circular dichroism (CD) spectroscopy. The CD spectrum of the variant showed only slight deviations from that of the WT Sfβgly (Fig. S5). Between 218 nm and 250 nm, the two spectra were nearly superimposable, whereas the largest differences occurred at 195 nm, where the variant displayed higher ellipticity, and at 207 nm, where it showed lower ellipticity than the WT. These features indicate that the overall topology is largely conserved despite modest changes in the secondary structure content. Thermal unfolding monitored by CD (**Figure 3B**) revealed a substantial loss of stability in the engineered enzyme, with a melting temperature (*T*_m_) of 44.0 ± 0.5 °C compared to 57.1 ± 0.4 °C for the WT. This reduction suggests that the introduced mutations disrupted secondary-structure packing, rendering the protein less stable. It is important to note that because this variant was not subject to any experimental screening, a decrease in thermal stability was anticipated; if thermostability had been a priority, initial screening of multiple PROSS-derived variants might have yielded more stable designs. The CD profile of WT Sfβgly closely matches that reported by Souza et al. 2016^12^, although the *T*_m_ measured here is 14 °C higher than previously reported. This discrepancy likely reflects differences in experimental conditions, including protein concentration, purification and buffer composition.

### Structure and binding of Pnp’s in the β-/α-variant

To investigate how the β-/α-variant accommodates the new stereoisomer while retaining activity its native substrates, each substrate used in the kinetic assays was docked into the AlphaFold^20,21^ predicted structure of the variant (**Figure 4** and Fig. S6). Except for Pnp-β-gal, all ligands produced lowest-energy docking models in which the sugar moiety occupied the –1 site of the catalytic pocket and the phenolic group occupied the +1 subsite, consistent with productive binding. Docking scores showed a strong correlation with experimental *k*_cat_ values for the β-/α-variant (**Figure 4B**). The relationship between docking score and catalytic efficiency was more moderate (*r* = –0.570, data not shown), because Pnp-β-fuc exhibited the highest *k*_cat_/*K*_M_ despite not having the most favorable binding score.

**Figure 4.**
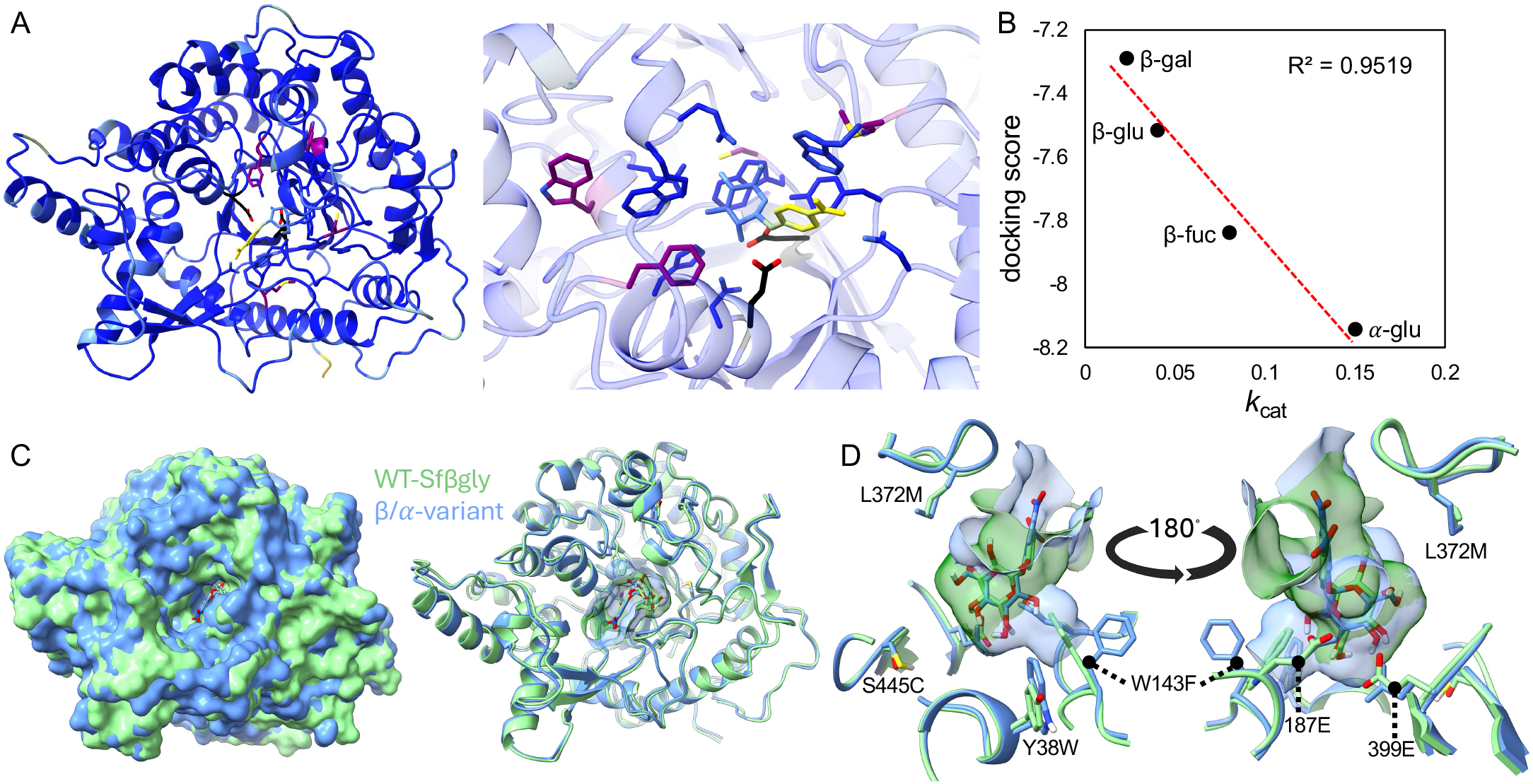
Structural modeling and substrate docking in the predicted β-/α-variant. (**A**) AlphaFold3 structure of the colored by pLDDT, predicted with Pnp-α-glu in the active site. Mutated residues relative to the WT are shown in purple, catalytic residues (E187 and E399) in black, and first-shell substrate-contacting residues in blue. (**B**) Correlation between docking score and experimentally measured *k*_cat_ values for each substrate. Docking scores correspond to the lowest-energy models generated using AutoDock Vina; full docking models are shown in Fig. S6. (**C**) Structural comparison of WT Sfβgly (green) docked with Pnp-β-glu and β-/α-variant (blue) docked with Pnp-α-glu. (**D**) Surface view of the catalytic pocket showing docked substrate. The W143F mutation in the variant (blue surface) creates additional space that facilitates accommodation of the α-configured substrate.

Structural comparison between the crystal structure of WT Sfβgly and the AlphaFold2-predicted structure of the β-/α-variant revealed a high degree of similarity (**Figure 4B** and Table S1), including the positioning of the catalytic residues E187 and E399 and three of the mutants introduced by FuncLib (Y38W, L372M, and S445C, Fig. S6C). Among all substitutions, the W143F mutation seems to contribute the most for the newly acquired α-activity. This mutation replaces a bulky tryptophan with a smaller phenylalanine, and its altered rotameric orientation creates additional space within the active site. This structural change enables Pnp-α-glu to bind in a flipped orientation relative to Pnp-β-glu in the WT enzyme (**Figure 4C-D**). Visual inspection of the docked complexes snows that the β-/α-variant accommodates the 2C hydroxyl of Pnp-α-glu (blue model, **Figure 4D**), allowing its anomeric carbon (1C) to occupy the same position as the 1C of Pnp-β-glu in the WT Sfβgly (green model). This offers a molecular rationale for how the engineered variant achieves α-hydrolytic activity. Overall, the structural predictions and docking analyses align well the experimental data and offer mechanistic insight into the emergence of dual β-/α-hydrolytic specificity in the designed glycosidase.

## Discussion

Protein design methodologies and structural prediction algorithms have advanced substantially in recent years, enabling increasingly precise manipulation of enzyme structure and function^22^. Despite notable success, engineering new enzymatic activities remains challenging due to constraints on functional encodability, the substantial time and resource investment required, and the frequent absence of efficient screening strategies^23^. In this study, I report the computational design of a β-glycosidase variant capable of hydrolyzing both β- and α-configured substrates, a rare within GH1, and without experimental screening. The resulting enzyme represents a one-shot design, demonstrating that targeted computational approaches can directly yield functional diversification and providing a proof of concept for dual β/α specificity in glycoside hydrolases.

Most reported engineering efforts in glycosidases have focused on enhancing native activities, such as hydrolysis or transglycosylation, improving properties, such as glucose tolerance and stability, typically through combinations of rational design, directed evolution, and computational methods^24,25^. Leveraging the inherent versatility of the (β/α)_8_ TIM-barrel fold which supports a wide range of catalytic functions^26^, the β-/α-variant described here extends this paradigm by enabling access to a non-native stereochemical configuration. This is important specially because in the literature, reported cases of GH processing both α- and β-configured substrates are rare, and no GH family has been classified with intrinsic dual α/β specificity for the same sugar.

An important mechanistic question is raised by this work concerns the catalytic pathway used for α-substrate hydrolysis. Is the mechanism an inverting or retaining for the new substrate Pnp-α-glu? Given that the engineered variant retains activity toward β-substrates, it suggests that the canonical retaining double-displacement mechanism remains operative. Dual α/β specificity within classical direct-displacement or double-displacement mechanisms proceeding through an oxocarbenium-like transition state has scarcely been documented^27–29^. The absence of clear precedents suggests that low-level α-activity in β-glycosidases may have been overlooked in prior studies due to detection limits or just lack of substrate. Although there are notable examples of GHs being able to deal with distinct stereochemistry. Dual stereochemical activity has been documented in GH109 and GH4 enzymes, which rely on NAD^+^-dependent redox chemistry compatible with both α-retaining and β-inverting reaction^30,31^, as well as in GH97 members that support both inverting and retaining mechanisms but remain restricted to α-substrates^7^.

From a design perspective, certain GHs appear inherently robust to functional diversification. Computational and experimental strategies have been successfully applied to glycosidases expand substrate utilization^32^ or to introduce dual activities, as demonstrated by combinatorial modular assembly of TIM barrel β/α repeats yielding activity toward both cellobiose and xylobiose^33^. In the present work, functional expansion was achieved from mutations in second-shell residues, which subtly reshaped the active-site environment. This strategy is consistent with previous observations that second-shell positions play a critical role in modulating substrate specificity while preserving catalytic function^16^. However, acquisition of the new function incurred a stability penalty, consistent with theoretical studies identifying second-shell residues as hotspots for activity–stability trade-offs^34^.

### Limitations of the study

The β-/α-variant exhibits lower expression levels than the WT Sfβgly, as observed during parallel purification from equivalent culture volumes (Fig. S2-S3). Two potential sources of expression inequality compared to the WT may lies in differences of plasmid and the codon sequence from the variant. Additionally, although α-hydrolytic activity was clearly detectable, turnover rates for Pnp-α-glu remain low, necessitating extended kinetic measurements over several hours. These limitations underscore the need for further optimization if the enzyme is to be applied to address specific functional or practical challenges.

## Resource Availability

### Lead Contact

Further information and requests for reagents and resources should be directed to and will be fulfilled by the lead contact, Felipe Otsuka.

### Material Availability

All materials generated in this study are available from the lead contact upon request and completion of an MTA.

### Data and Code Availability

Any additional information required to reanalyze the data reported in this paper will be shared by the lead contact upon request. This paper does not report original code.

## Acknowledgements

I am grateful to Sandro R. Marana for providing the plasmid containing the wild-type construct. I also thank Sandro R. Marana and Ingemar André for granting me access to their laboratories to produce and analyze the enzymes.

This work was supported by FAPESP (Fundação de Amparo à Pesquisa do Estado de São Paulo; Grants 2016/22365-9, 2018/18537-4 and 2018/25952-8).

## Author Contributions

Writing—original draft: F.A.M.O.

Writing—review & editing: F.A.M.O.

Conceptualization: F.A.M.O.

Investigation: F.A.M.O.

Funding Acquisition: F.A.M.O.

## Declaration of generative AI and AI-assisted technologies in the writing process

During the preparation of this work the author used Copilot from Microsoft to only review grammar and to improve readability. After using this tool, the author reviewed and edited the content as needed and takes full responsibility for the content of the published article.

## Methods

### Reagents, plasmids and buffers

The WT Sfβgly is inserted in a vector pET-46 EK-LIC (Merck Millipore, Billerica, MA, USA).23 The β-/α-variant was purchased cloned in pET-28a(+) by GenScript. Broad range SDS-PAGE standards were purchased from Bio-Rad (Hercules, CA, USA). F100 buffer stands for 100 mM potassium phosphate buffer, pH6; lysis binding buffer is 100 mM sodium phosphate containing 300 mM NaCl, 20 mM imidazole and glycerol 12%, pH 7.45; elution buffer is 100 mM sodium phosphate containing 300 mM NaCl, 300 mM imidazole and glycerol 5%, pH 7.45. Isopropyl β– D–1–thiogalactopyranoside (IPTG). 4-Nitrophenyl β-D-glucopyranoside, 4-Nitrophenyl β-D-galactopyranoside, 4-Nitrophenyl β-D-fucopyranoside and 4-Nitrophenyl α-D-glucopyranoside from fisher scientific.

### Designing of β-/α-variant and Pnp docking

The X-ray structure of the Sfβgly, (PDB id: 5CG0), chain A and B, the dimer, was used as input to Rosetta FastRelax^14^. Then the minimized structure was submitted in PROSS website “https://pross.weizmann.ac.il/step/pross-terms/"^15^ with restriction to not modify 42 positions involved in the binding and catalysis of substrates and in the interface of the dimer. Selection of the model was based on observation of the models using PyMOL and the suggestions, such as avoiding sequences with more than 10 mutated residues in segments. The selected PROSS model was then directly submitted to FuncLib^16^ with following parameters: Amino acid positions to diversify: 38A, 143A, 372A, 400A, 445A; Essential amino acid residues: 399A, 187A, 97A, 331A, 329A, 39A, 142A, 186A, 444A, 451A, 452A; Ligand to keep during simulations: none; Ions to keep during simulations: none. The only model with tolerable sequence space was selected to test in experiments.

Docking analysis started with the structure of the .sdf format of 3D structure of ligands downloaded from the PubChem, the X-ray structure of the WT Sfβgly and the Alpha Fold2 predicted structure of the β-/α-variant. The docking program used was AutoDock-Vina^35^ version 1.2 and each ligand was docked individually in the protein models using exhaustiveness set at 124. Score analysis was based on the output from Vina, using the best score from the model with the substrate correctly inserted in the active site, *i*.*e*., the sugar part in the –1 site of the catalytic pocket.

### Expression and purification of the glycosidases

Both the Sfβgly and variant were confirmed by sequencing, expressed and purified in parallel. Their vectors pET46-Sfβgly and pET-28a(+) were separately transformed into chemically competent *E. coli* BL21 (DE3) and a single colony was inoculated in 5 mL of LB to make a pre-inoculum with antibiotics, 50 µg/mL carbenicillin for pET46-Sfβgly and kanamycin pET-28a(+), and incubated under shaking at 200 rpm, 35 °C for 16 h. Following that, 1 mL of this pre-inoculum was added into 500 mL of LB with antibiotics. The inoculum was incubated at 37 °C under shaking (200 rpm) until it reached OD600 of 0.5 – 0.7. After that IPTG was added to a final concentration of 0.8 mM to induce the expression of both glycosidases for 16 h at 25 °C and shaking at 200 rpm. Then, cells were harvested through centrifugation at 5,000 g for 20 min at 4 °C and stored at –20 °C for further processing.

The pelleted cells were suspended in lysis binding buffer (2:1 w/v) containing lysozyme (1 mg/mL). Cell lysis was done by sonication (Branson Sonifier 250, Soni-tech) using 5 pulses at 40% amplitude for 10 s with 20 seconds interval between pulses. Samples were kept in the ice during the sonication. Cell debris were sedimented by centrifugation at 9,000 g (30 min, 4 °C). The supernatant was injected in a column FF-crude Ni-NTA resin (Cytiva) which was pre-equilibrated with lysis binding buffer using an ÄKTAFPLC (GE HealthCare). The elution used an protocol to wash weakly bound proteins with 2 CVs of binding buffer, followed by a linear gradient with elution buffer with 5 CVs. Fractions were collected in 96 deep well plates and those containing Sfβgly were pooled and buffer exchanged to F100 in a centrifugal filter device (10 kDa cutoff; Amicon Millipore) with repetitive cycles of centrifugation and washing with F100 buffer. Purification evaluation was made by SDS-PAGE.44 A Nanodrop was used to quantify the protein concentration based on the extinction coefficient calculated by ProtParam, WT Sfβgly = 120,460 and variant = 128,480.

### Size exclusion chromatography (SEC)

The SEC was performed using an ÄKTAFPLC. A Superdex 200 Increase 10/300 GL (GE HealthCare) column was pre-equilibrated with buffer F100. Concentrated glycosidases were loaded into the columns using a 200 µL sample loop. Chromatography data were collected and processed using the UNICORN software version 7.12 (Cytiva).

### Enzymatic assay

The initial rate (*v*_0_) of hydrolysis of at least nine different concentrations of each 4-nitrophenol substrate was prepared in F100 buffer and used to determine the kinetic parameters (kcat and Km) assuming a steady-state kinetics. In a 96-microtiter plate sat on ice each substrate were aliquoted (50 µL/well) in a combination of duplicates, different concentrations and repeating for four different time points. Then, 50 µL of purified enzyme in F100 was added in each well to a final concentration of 1 mM. Enzymatic reaction started when the assay plates were transferred from ice to a warm bath at 30 °C. In order to determine the initial rate, reactions for each substrate concentration were performed along four times intervals, for Pnp-α-glu: 15, 35, 55 and 75 min, for Pnp-β-glu and Pnp-β-fuc: 5, 10, 15 and 20 min, for Pnp-β-gal: 30, 70, 115 and 170 min. Reactions were stopped by addition of 0.5 M Na2CO3 (100 µL). Absorbance at 415 nm was measured in a microplate reader Elx800 (Biotek) and converted to mols of the released product (4-nitrophenol) using a linear calibration curve. Product (nmols) was plotted versus time intervals and initial reaction rate (*v*_0_ nmols/s) determined based on the linear coefficient. Only linear plots were used. The concentration of the substrate and respective *v*_0_ were fitted in the Michaelis-Menten equation to obtain the kinetic parameters (Km, kcat and kcat /Km). Assays for kinetic parameters determination were made in duplicates.

### Circular Dichroism (CD) and melting temperature (*T*_m_)

CD was carried out in a Jasco J-815 CD spectropolarimeter (Jasco Scandinavia AB) attached to a Peltier system. Data were collected using a rectangular quartz cuvette (NSG Precision Cells Inc., Farmingdale, NY, USA) with a path length of 0.2 cm in continuous scanning mode at a speed of 20 nm·min^−1^, an integration time of 1 s, a bandwidth of 1 nm and standard sensibility. The samples were 300 μL of purified enzyme in F100, at concentrations of 35 μg/mL for the WT Sfβgly (0.6 μM) and 60 μg/mL for the β-/α-variant (0.9 μM). Two spectra were collected from 192 to 260 nm at 20 °C to 90 °C and averaged. Molar absorptivity (Δε) was calculated using the BestSel^36–38^. Estimation of the protein *T*_m_ was made using a python script generated with Microsoft Copilot that fits the experimental data of ellipticity measured at 222 nm to a two-state Boltzmann model:

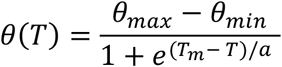

where *θ*(*T*) is the ellipticity at temperature *T, θ*_max_ and *θ*_min_ represent the ellipticity values of the unfolded and folded states, respectively, *T*_m_ is the melting temperature, and *a* corresponds to the slope factor of the linear region of the sigmoidal transition.

