## Supplemental figures for "Engineering a bifunctional alfa and beta hydrolase from a GH1 beta-glycosidase"

Figure S1

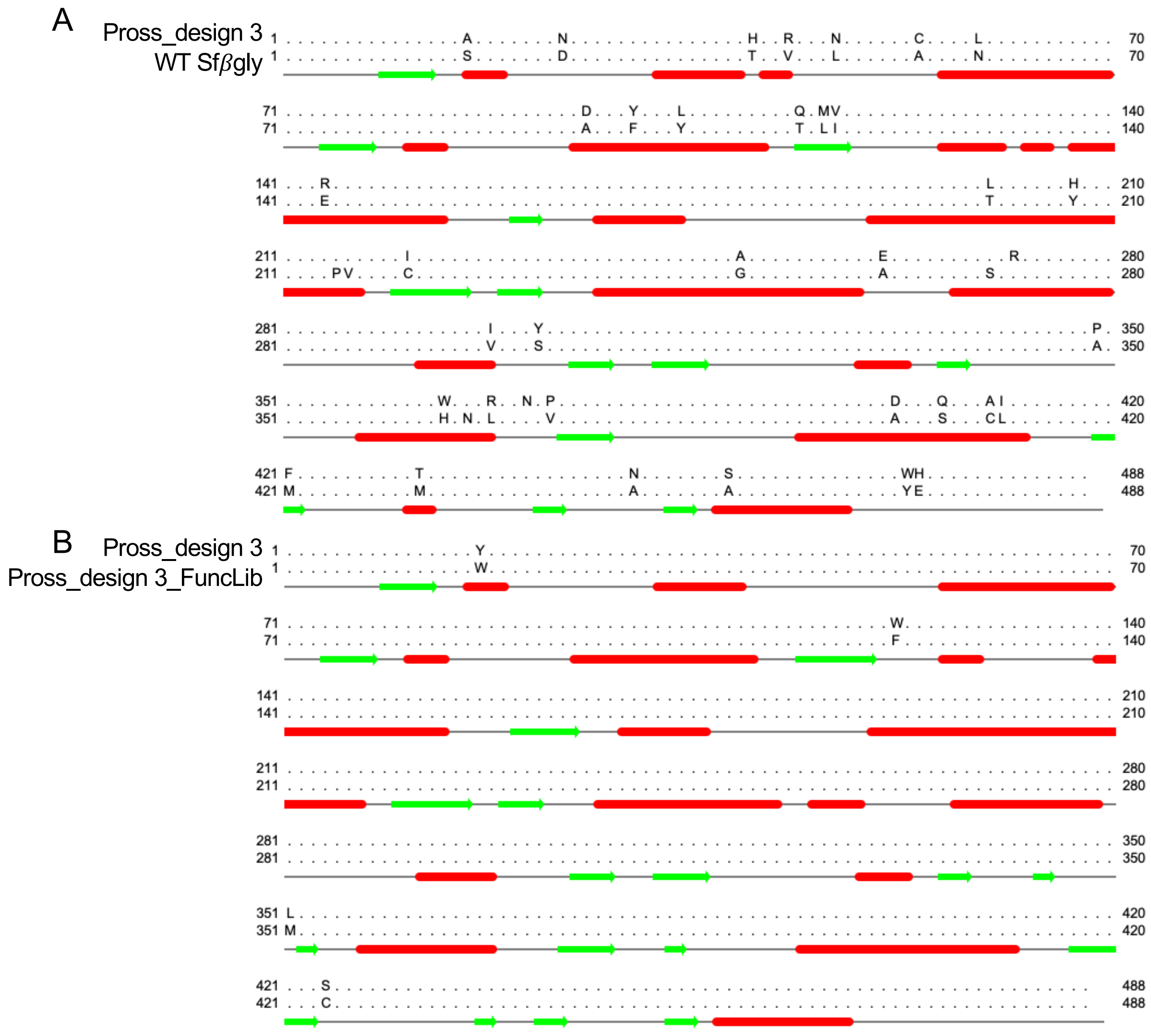

Figure S1. Alignmnet between (A) the first design (PROSS_design3) with the WT Sfβgly containing the secondary structure from the PDB id 5CG0. In (B) the alignment between the first mutant variant with the variant choose for the experimental assays, also addressed a β-/α- variant containing its secondary structure from the AlphaFold predicted structure. Aligned dots indicate the same amino acid. Residue counting is shifted 21 amino acids less, thus residue 1 is equivalent to 22 when describing positions. Figure generated with Jalview.

Figure S2

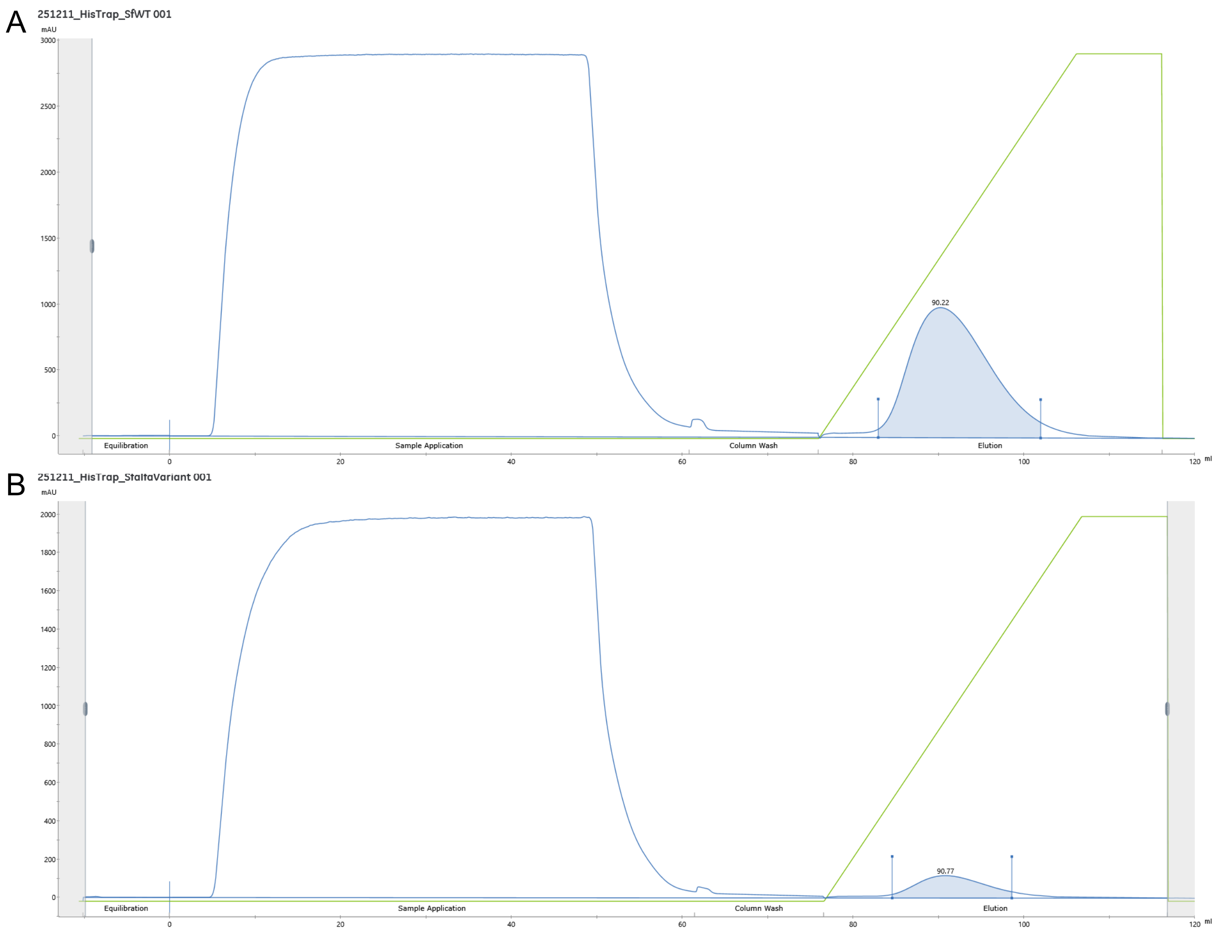


Figure S2. Purification of WT Sfβgly (**A**) and β-/α-variant (**B**). Sample injection and elution in HisTrap™ ff-crude column. Blue line is absorbance at 280 nm and green line is the concentration of the elution buffer over the gradient.

Figure S3

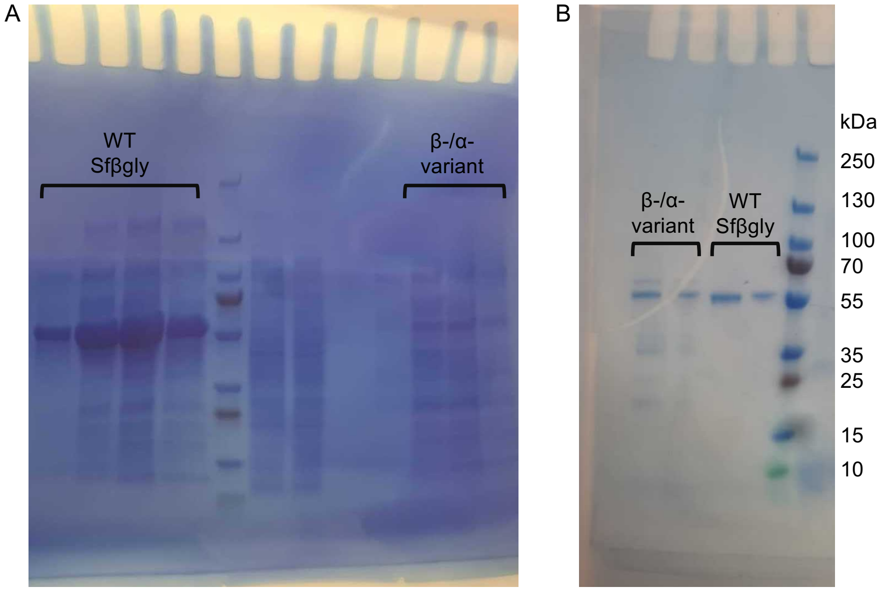


Figure S3. Purified β-/α-variant and WT Sfβgly. (**A**) Elution during Ni-NTA resin in ff-crude column. (**B**) Pooled fractions of Size-exclusion chromatography after Ni-NTA purification.

Figure S4

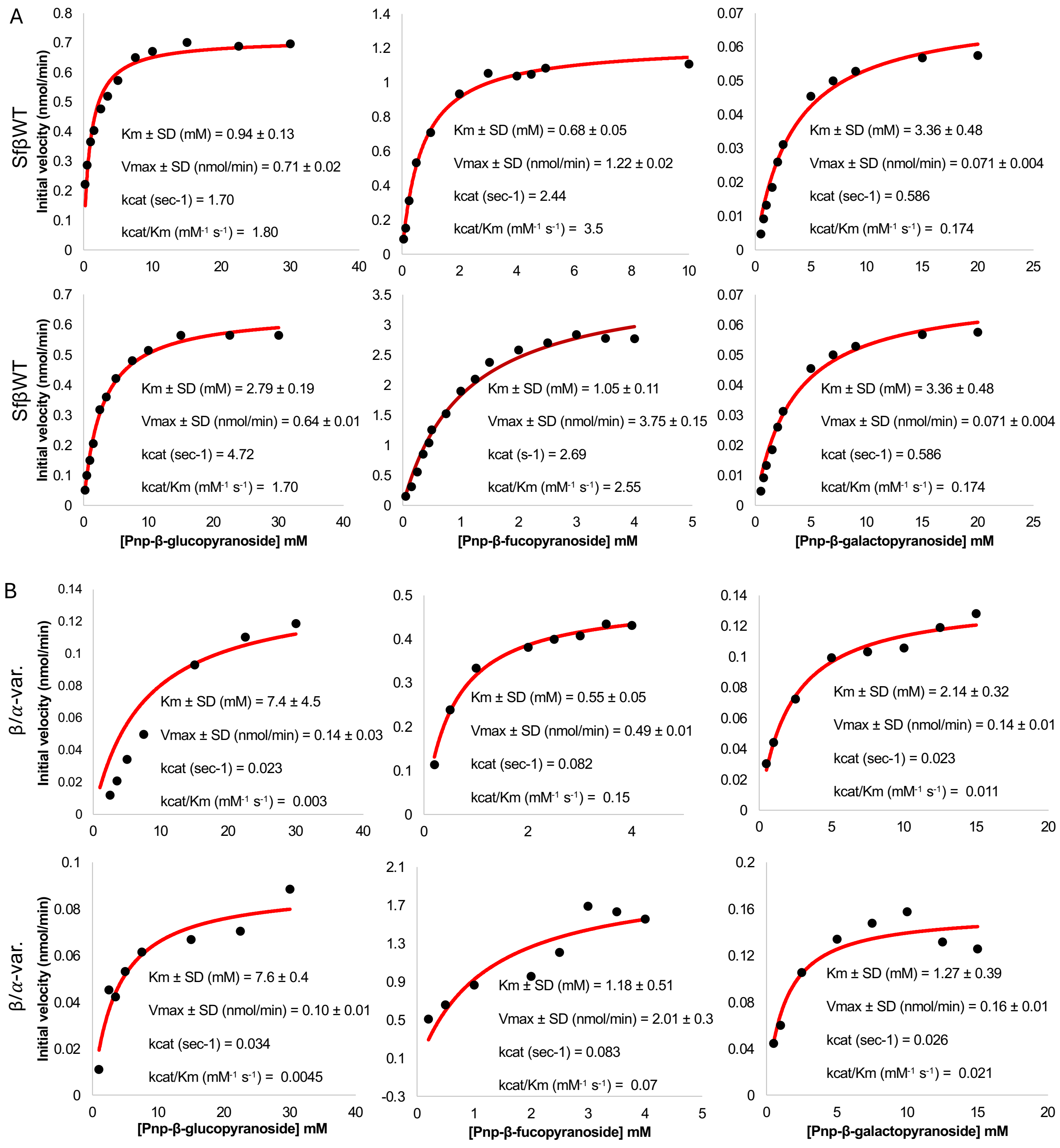


Figure S4. Enzymatic assays of WT Sfβgly (**A**) and β-/α- variant (**B**). Fitted Michaelis-Menten equation is in red and experimental slopes as black dots.

Figure S5

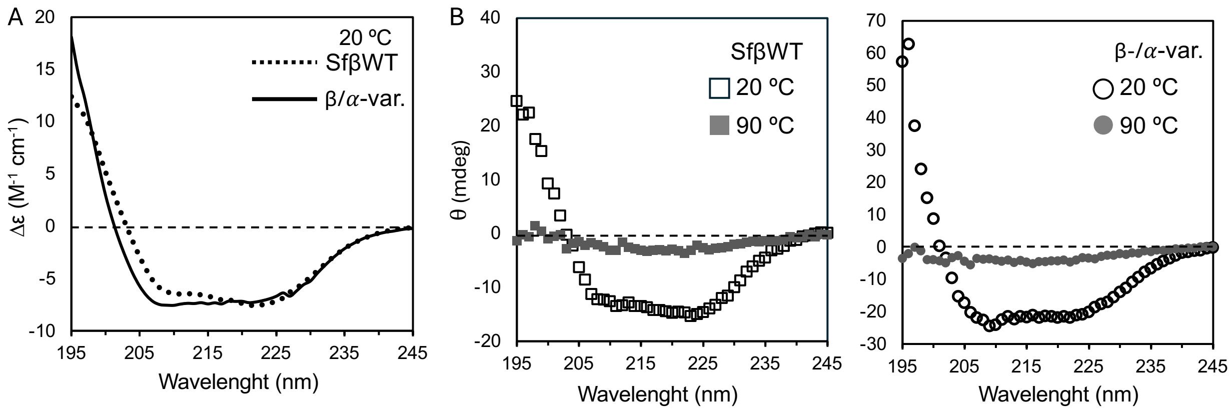


Figure S5. Circular dichroism profile of β-/α-variant and WT Sfβgly. (**A**) Lines (solid and dotted) represent fitted values of molar absorptivity calculated in the BeStSel website. (**B**) CD profile of β-/α-variant and WT Sfβgly in two temperatures, in native condition (20 °C) and unfolding condition (90 °C). The average of three measurements are shown.

Figure S6

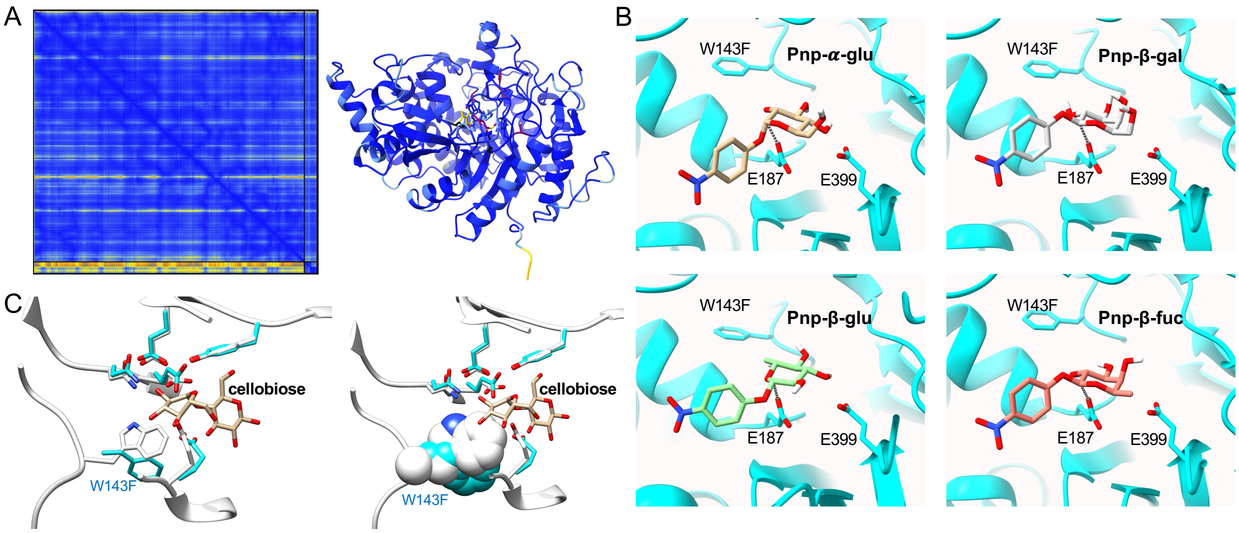


Figure S6. Models and docking of ligands in the β-/α-variant. (**A**) AlphaFold3 predicted aligned error of the β-/α-variant with Pnp-α-glu. (**B**) Docking models generated with Vina using the structure predicted with AlphaFold2. Distance (grey pseudobond) between C1 and O from E187 in each Pnp's are: Pnp-α-glu = 4.02 Å, Pnp-β-glu = 4.3 Å, Pnp-β-fuc = 4.4 Å, Pnp-β-gal = 4.2 Å. (**C**) Docking of cellobiose in the variant. Variant model is colored in blue or cyan. Model in white correspond to the WT Sfβgly. Note the non-reducing end of the sugar is in the –1 site as standard in hydrolysis.

Table S1. Comparison of β-/α- variant and WT Sfβgly constructs in plasmid for expression calculated with protparam.

| Enzyme | n° a.a. | Mw (g/mol) | Theoretical PI | Negatively charged residues | Positively charged residues |
|---|---|---|---|---|---|
| WT Sfβgly | 504 | 57,950 | 4.8 | 78 | 45 |
| β-/α- variant | 509 | 59,014 | 5.3 | 74 | 51 |

Ordered variant is a construct containing a $His_6$-tag in the N-terminal followed by a thrombin cleavage site (MGSSHHHHHHSSGLVPRGSH). The WT construct N-terminal contains an enterokinase cleavage site after the $His_6$-tag (MAHHHHHHVDDDDKI).
